# A new full-length virus genome sequencing method reveals that antiviral RNAi changes geminivirus populations in field-grown cassava

**DOI:** 10.1101/168724

**Authors:** Devang Mehta, Matthias Hirsch-Hoffmann, Mariam Were, Andrea Patrignani, Hassan Were, Wilhelm Gruissem, Hervé Vanderschuren

## Abstract

Deep-sequencing of virus isolates using short-read sequencing technologies is problematic since viruses are often present in complexes sharing a high-degree of sequence identity. The full-length genomes of such highly-similar viruses cannot be assembled accurately from short sequencing reads. We present a new method, CIDER-Seq (Circular DNA Enrichment Sequencing) which successfully generates accurate full-length virus genomes from individual sequencing reads with no sequence assembly required. CIDER-Seq operates by combining a PCR-free, circular DNA enrichment protocol with Single Molecule Real Time sequencing and a new sequence deconcatenation algorithm. We apply our technique to produce more than 1,200 full-length, highly accurate geminivirus genomes from RNAi-transgenic and control plants in a field trial in Kenya. Using CIDER-Seq we can demonstrate for the first time that the expression of antiviral doublestranded RNA (dsRNA) in transgenic plants causes a consistent shift in virus populations towards species sharing low homology to the transgene derived dsRNA. Our results show that CIDER-seq is a powerful, cost-effective tool for accurately sequencing circular DNA viruses, with future applications in deep-sequencing other forms of circular DNA such as transposons and plasmids.

## INTRODUCTION

Advances in high-throughput sequencing technologies have produced extensive data about viral diversity, both at the species/isolate level and at higher taxonomic levels. The increasing use and applications of high-throughput sequencing technologies have also revolutionized virology by identifying and characterizing previously unknown viruses. The abundance of new virus sequence data recently led to the publication of a consensus statement proposing revisions in virus taxonomy incorporating metagenomic sequence data (Simmonds et al. 2017). However, the contribution of high-throughput sequencing technologies to virus detection and identification remains constrained by several technical challenges including the risk of assembling artificially chimeric viral genomes due to short sequencing read lengths.

Sequence-bias during virus enrichment before sequencing is also a recognized drawback during virus deep-sequencing. Amplification methods to enrich viral nucleic acids rely on using either Polymerase Chain Reaction (PCR) with primers designed to bind a conserved sequence in the genome, or random circular amplification (RCA) utilizing the unique properties of the Phi29 DNA polymerase coupled to random-nucleotide primers (Dean et al. 2001). PCR-based methods have an obvious drawback because differing primer-template affinities can result in a loss of viral templates (Sipos et al. 2010). RCA relies on random primers and therefore is less prone to primer complementarity bias. However, RCA results in hyper-branched, high molecular weight, concatenated products (Lasken and Stockwell 2007) that must be linearized through the use of restriction enzymes (REs) (Inoue-Nagata et al. 2004) or mechanical shearing prior to Sanger or NGS sequencing. The use of REs necessarily requires prior information on conserved RE sites and therefore results in the loss of viral sequences that may have none, or multiple recognition sites for the specific REs used. Another approach called polymerase cloning or “ploning” (Zhang et al. 2006) employs endonucleases and DNA repair enzymes to linearize RCA products, followed by shotgun sequencing and whole genome assembly.

RNA interference (RNAi), i.e. the production of short interfering RNA (siRNA) from double stranded RNA (dsRNA) substrates to induce gene silencing, has been effectively used to engineer virus resistance to RNA and DNA viruses in a number of crop plants (Rey and Vanderschuren 2017; Pooggin 2017). Field assessment of RNAi-mediated resistance against *Bean golden mosaic virus* (BGMV) in transgenic common bean (*Phaseolus vulgaris*) (Aragão and Faria 2009) and against *Tomato yellow leaf curl virus* (TYLCV) in transgenic tomato (*Solanum lycopersicum*) (Fuentes et al. 2016) indicate that the RNAi-mediated genetically engineered resistance to DNA viruses is stable under field conditions when plants are exposed to single viral species. However, the effectiveness of RNAi technology in crops suffering from infections by multiple virus species has not yet been assessed. Because RNAi technology depends on sequence complementarity between the transgene and the targeted viral sequences, it is essential to determine the degree of sequence complementarity required for broad spectrum virus resistance in the field.

Here we report the development of a new technique for sequence-independent viral DNA enrichment utilizing assembly-free Single Molecule Real Time (SMRT; Pacific Biosciences Inc.) long-read sequencing called CIDER-Seq (<u>C</u>ircular <u>D</u>NA <u>E</u>nrichment <u>s</u>equencing). CIDER-Seq requires only an estimate of virus genome size and, optionally, a single reference sequence. We have also developed a new algorithm, DeConcat, to parse concatenated DNA strands that are generated by rolling circle amplification (RCA), allowing us to sequence complete geminivirus genomes without reference-based or *de novo* assembly. We demonstrate the effectiveness of CIDER-Seq by producing 1328 full-length genomes of cassava mosaic geminiviruses (CMGs). CMGs cause cassava mosaic disease (CMD) and severe economic losses for farmers, particularly in sub-Saharan Africa (Rey and Vanderschuren 2017). To date, nine CMG species have been identified that share 68% to 90% sequence identity based on Sanger sequencing of PCR and RCA products (Rey and Vanderschuren 2017). CMGs are whitefly-transmitted, bipartite viruses of the genus *Begomovirus* in the family *Geminiviridae*—the most populous family of eukaryote-infecting viruses (International Committee on Taxonomy of Viruses 2016). They are comprised of two separate genomes, designated DNA A and DNA B (Rey and Vanderschuren 2017). Our CIDER-Seq enabled analysis of CMG populations in field-grown transgenic cassava plants and provided new insights into the limits of RNAi mediated virus resistance.

## RESULTS

In this study we profiled virus populations from a field trial of previously developed transgenic (Vanderschuren et al. 2009) and control cassava lines, conducted in Kenya. The transgenic cassava lines (dsAC1-101 and dsAC1-152) produce 155bp long dsRNA targeting the conserved 3’-end of the *AC1* gene (located on DNA A) from African cassava mosaic virus (ACMV) species (Vanderschuren et al. 2009). The field trial was conducted in 3 replicated randomized blocks over a span of 30 weeks. Disease symptom, weather and whitefly count data was recorded weekly, starting at 8 weeks after planting (Additional File 1, Supplementary Fig. 1 a–e). Transgenic plants showed no symptoms until week 17, although by the end of the experiment these lines showed between 60–70% mean disease incidence (Additional File 1, Supplementary Fig. 1a, b). Leaf samples were collected at the completion of the field trial and assessed for virus levels (Additional File 1, Supplementary Fig. 1f) and sequence diversity.

Using leaf samples from cassava plants with CMD symptoms we first enriched complete DNA genomes of CMGs (Fig. 1) by automated size selection of DNA molecules in the 2.8 kb size range (Step 1). Phi29 DNA polymerase RCA of the selected DNAs was carried out using random hexameric primers (Step 2). Using a ‘ploning’(Zhang et al. 2006) based protocol, we amplified the products of the initial amplification in an additional primer-free Phi29 DNA polymerase reaction to further “de-branch” the hyper-branched DNA (Step 3). Next, the hyperbranched structure was resolved using ssDNA-digesting S1 Nuclease (Step 4) and repaired with DNA Polymerase I and T4 DNA Polymerase (Step 5). The linear DNA fragments were purified using magnetic beads (Step 6), which excluded small DNA fragments produced during the RCA steps. Semi-quantitative PCR was used to assess the relative importance of each successive step in the enrichment protocol (Additional File 1, Supplementary Fig. 2).

**Figure 1:**
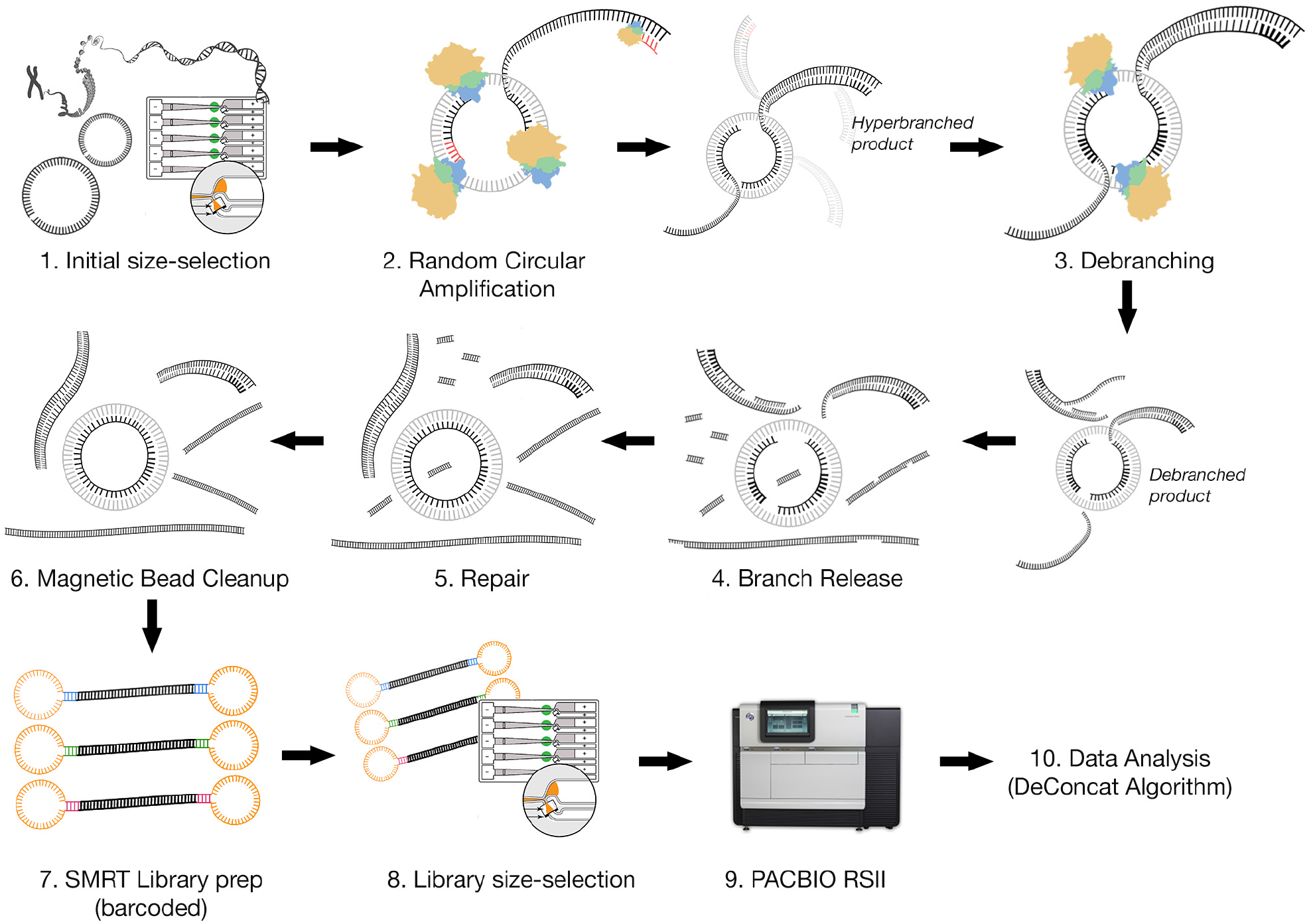
Enrichment of circular DNA based on automated size selection, non-denaturing random circular amplification (RCA), linearization and repair of the RCA product followed by Single Molecule Real Time (SMRT) library creation.

The enriched samples were next barcoded and pooled to build 4 separate SMRT-sequencing libraries (Additional File 1, Supplementary Table 1). SMRT Cell 1 (and 4) were used primarily to refine the CIDER-Seq and DeConcat methods and Cells 2 and 3 were used for analyzing virus populations in transgenic and control plant lines. The raw sequencing reads were next filtered for high-quality insert sequences (predicted minimum quality of 99.9 and minimum insert length of 3000bp). As expected, a majority of ROIs were greater than the 3 kb size cut-off, indicating that the inserts were comprised of concatenated RCA products.

We developed a custom data analysis pipeline to resolve the concatenated sequences into their component parts. First, in a basic filtering step all ROIs were binned into non-viral, CMG DNA A and CMG DNA B sequences. The results from this step indicated that at least 98% of the ROIs in all libraries were found in the CMG bins (Fig. 2). Next, we trimmed non-viral DNA from the ends of each ROI by performing a circular DNA-aware multiple sequence alignment (see Methods).

**Figure 2:**
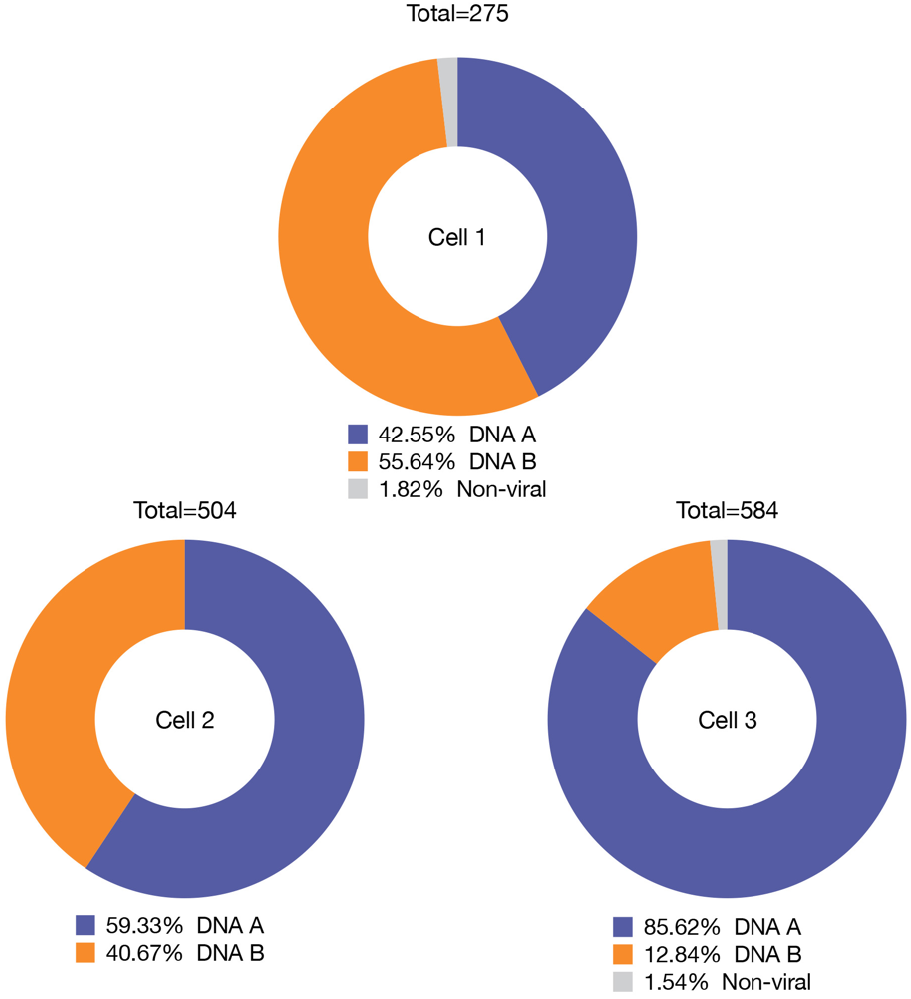
Proportion of viral (DNA A & DNA B) and non-viral reads produced in each SMRT Cell.

These trimmed ROIs were next passed through the de-concatenation algorithm we termed ‘DeConcat’ (Fig. 3). DeConcat begins (Step 1, Fig. 3) by cleaving the trimmed ROI (A-B’) at the 30nt position to produce two segments (A-A’ and B-B’) and aligning them using MUSCLE (Step 2a). (This fixed distance of 30 nt was determined by benchmarking a range of values from 500 nt to 30 nt. The benchmarking results are shown in Supplementary Fig. 3) Using the alignment consensus, a score is calculated by dividing the total consensus length by the number of consensus fragments separated by gaps (Step 3). The algorithm iterates back (Step 4) to step 1, increases the cleavage position by 30, proceeds to step 2a and step 3 and iterates back to step 1. This proceeds until all possible cleavage positions have been used. The algorithm retains the alignment with the highest score in step 3. A second iteration now aligns the reverse complement sequence of the first segment (i.e. converts A-A’ to A’-A) (Step 2b), calculates the score, and if the score is higher than the previously retained alignment, the reverse complement alignment is used. If the computed score of the two best aligned segments is > 20, a 10nt sliding window on both ends is applied and the first windows with >90% identity are used to determine start and end positions of the alignment fragments. The final retained alignment for each ROI can take one of eight possible overlap patterns (Step 5, Fig. 3). If the smaller segment lies completely within the larger one (cases 1a-1d) the algorithm re-starts with the larger segment (Step 6) and eliminates the smaller one. If the second segment (B-B’) overlaps the end of the first segment (A-A’) (case 2), the algorithm restarts (step 6) with both segments as independent ROIs. If the B-B’ overlaps with the front of A-A’ (case 3), the overlapping part of B-B’ is eliminated and the remaining segment (B-B”) is reattached in its original position at the end of A-A’. If B-B’ overlaps the end of A’-A (i.e. the reverse complement of A-A’ produced in step 2b) (case 4), the overlapping part of A’-A is eliminated, the remaining segment (A’-A”) is reverse complemented and B-B’ is re-attached to the end of A”-A’. In case 5, B-B’ overlaps with the start of A’-A. Here, the overlapping part of B-B’ is eliminated to produce B-B”. A’-A reverts back to its original configuration A-A’ and the two segments (A-A’ & B-B”) are reattached. Once case-resolution (step 5) is completed, the resulting sequence is entered back into the algorithm at step 1 for another round of de-concatenation. The case resolutions have been designed so as to maintain the integrity of the initial fragment, i.e. the order in which bases are produced by the sequencer is never changed. Effectively, case-resolution simply results in sequence reduction from the ends.

**Figure 3:**
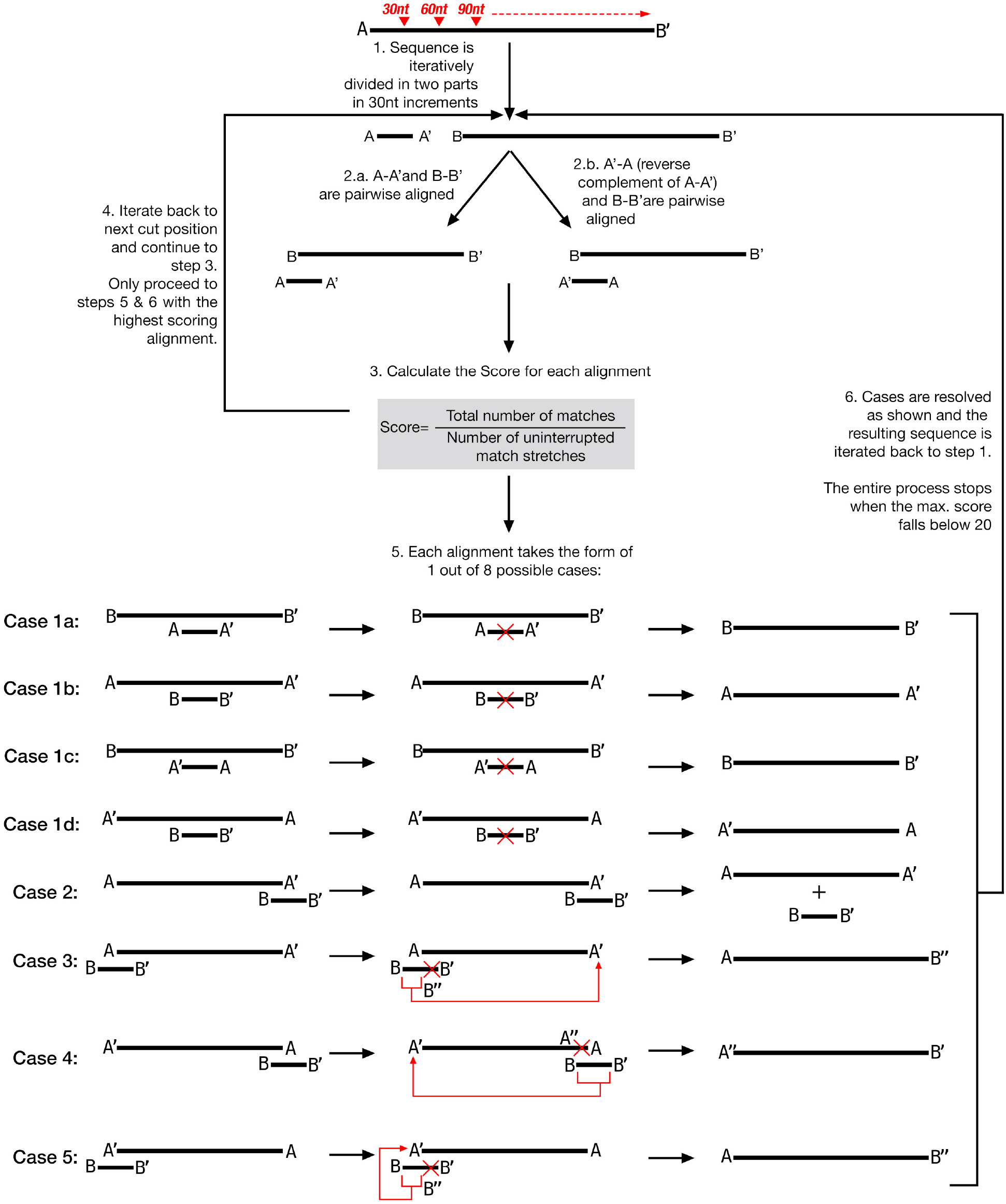
Flowchart for the DeConcat algorithm

DeConcat performance stats were derived by analyzing the data from SMRT Cells 1 and 4 (also known as SS for size selected and NSS for non-size selected respectively). After running DeConcat on, we found that in both NSS and SS data-sets a majority of sequences had scores in the range of 4–6 at the end of the DeConcat program (Additional File 1, Supplementary Fig. 4a), indicating that for these sequences the final de-concatenation round had indeed resolved most repeat sequences. Interestingly, most reads in the SS data set required just 1–3 rounds of deconcatenation to resolve their sequences (Additional File 1, Supplementary Fig. 4b). We also found that of the eight possible alignment cases in DeConcat, cases 1a, 1b and 3 were the most frequent (Additional File 1, Supplementary Fig. 4c) in both data sets. It should be noted however that cases are assigned by DeConcat iteratively and hence do not reflect overall sequence topology. It is thus likely that the frequency of cases 1a and 1b is simply due to alignments of fragments differing greatly in size during DeConcat rounds. The relatively high frequency of case 3, however, does suggest that RCA-derived sequences are often direct sequence concatemers. Overall, based on frequencies of cases 1c, 1d, 4 and 5 it appears that cases where concatemers are formed between sequences and their opposite strands are rare.

To test the hypothesis that more DeConcat processing rounds usually result in shorter sequences we plotted the number of DeConcat rounds per sequence against its length (Additional File 1, Supplementary Fig. 4d). However, in many cases, several processing rounds still resulted in sequences ~2800 bp in length—further demonstrating the ability of the system to resolve RCA products into single molecules. We also tested the ability of DeConcat to process RCA-derived long sequencing reads in the absence of a reference sequence that is used in the trimming step. When comparing results with and without the trimming step, we found that the lengths of only 8.8-15.3% of sequences were affected by omitting the trimming step. Further, the average change in length of these sequences was only 15-30 bp (Additional File 1, Supplementary Fig. 5). We conclude that DeConcat can effectively resolve RCA-derived sequences even in the absence of a reference sequence and does not require prior information on expected monomer length distributions. However, the trimming step is recommended to improve sequencing in cases where multiple viral molecules may be joined together due to the action of the Phi29 DNA polymerase.

The resulting de-concatenated viral ROIs had a greatly reduced size distribution (Fig. 4), with a clear peak at 2.8 kb in all three SMRT cells.

**Figure 4:**
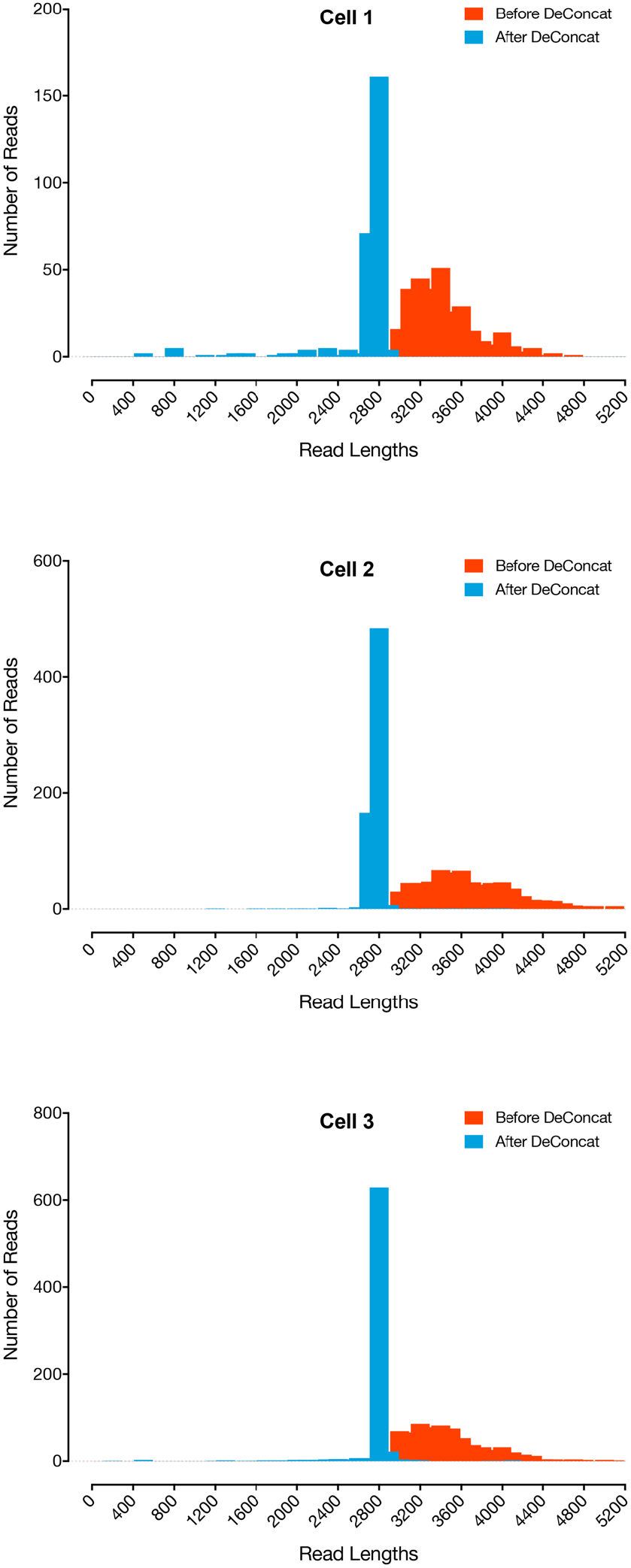
Size distribution of sequencing reads before (Red) and after (Blue) DeConcat processing per SMRT Cell.

Thus, DeConcat was able to reduce RCA-generated SMRT-Sequenced ROIs of a wide size-distribution into expected CMG-length virus sequences. This result validated the parameters and scoring formula applied in DeConcat and also demonstrated that the Phi29 DNA polymerase enrichment step amplified circular DNA in the 3kb range with high fidelity.

We performed a tBLASTn using the final DeConcat reads against virus protein reference sequences to annotate the virus genomes. This also allowed us to detect and classify sequencing errors in our final genomes (Additional File 1, Supplementary Fig. 6). The annotation results revealed frameshift mutations in one or more ORFs in several reads. Frameshift mutations are caused by insertions or deletions—the most frequent type of errors in SMRT sequencing (Laehnemann et al. 2016). To distinguish if these frameshifts were of biological origin or SMRT-Sequencing errors, we relaxed the minimum quality threshold of 99.9% for ROIs in the initial filtering step to 99.5% and 99%. When comparing the frequency and number of frameshift mutations between the three quality thresholds we found that the number of frameshifts per sequence increased with decreasing quality thresholds (Additional File 1, Supplementary Fig. 6a). and the frequency of frameshifts in each viral protein increased linearly with protein length (Additional File 1, Supplementary Fig. 6b), indicating that they are likely caused by SMRT sequencing errors. Between 5 to 8% of reads in the >99.99 quality threshold dataset have no frameshift errors (Additional File 1, Supplementary Fig. 6c).

We next performed a multiple sequence alignment of the full-length genomes against the reference virus sequence (ACMV-NOg isolate) from which the transgene was originally derived. This allowed us to assess the proportion of viruses in each plant line which can be efficiently targeted by the transgene. We found that the control plants had two populations of viruses, one with >90% identity to the reference ACMV-NOg and one with less than 75% (Fig. 5). In the transgenic lines, on the other hand, no sequences with greater than 90% identity to the target virus were detected and all the virus genomes belonged to the <75% identity category. This suggests that the transgenic plants could effectively limit target CMG species, leading to a surge in the proportion of non-target CMG species. Similar results were found upon analyzing only the virus sequences corresponding to the transgene region (Additional File 1, Supplementary Fig. 7).

**Figure 5:**
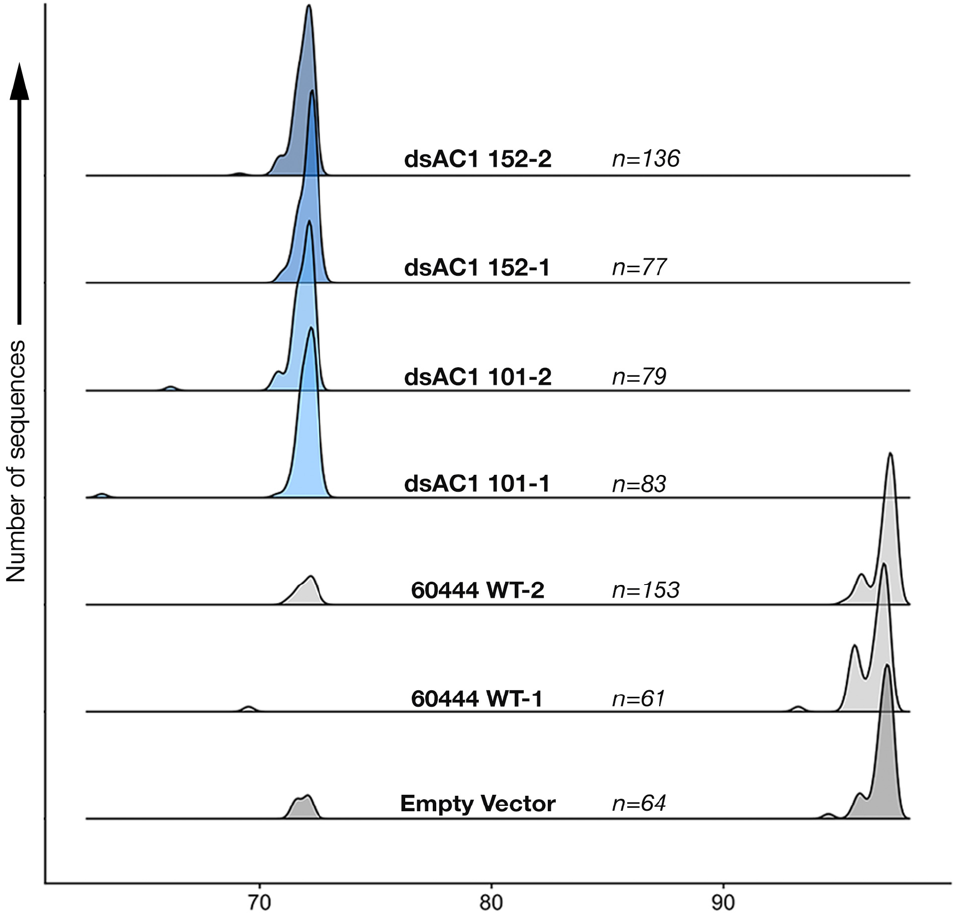
Density plot showing the proportion of virus sequences with their % identity to the ACMV-NOg genome from which the transgene was derived.

We next simulated the production of anti-viral 21nt siRNAs from each plant line. Putative siRNA sequences were aligned against each virus genomes and plotted according to the number of mismatches between the siRNA and the target virus. While, on average, between 75 and 100 transgene-derived siRNAs could target each virus per control sample with no mismatch, only between 10 and 15 siRNAs could target each virus in the transgenic lines with complete fidelity (Fig. 6). The presence of identical levels of predicted siRNAs with single mismatches in control and transgenic samples suggests that resistance mainly relies on perfectly matching siRNAs. Phylogenetic trees from all viruses in control and transgenic lines, along with reference genomes from all 7 CMG species found in Africa revealed that the non-target viruses identified in Fig. 5 are closely related to the *East African Cassava mosaic virus* and the *East African Cassava mosaic Cameroon virus* species (Fig. 7).

**Figure 6:**
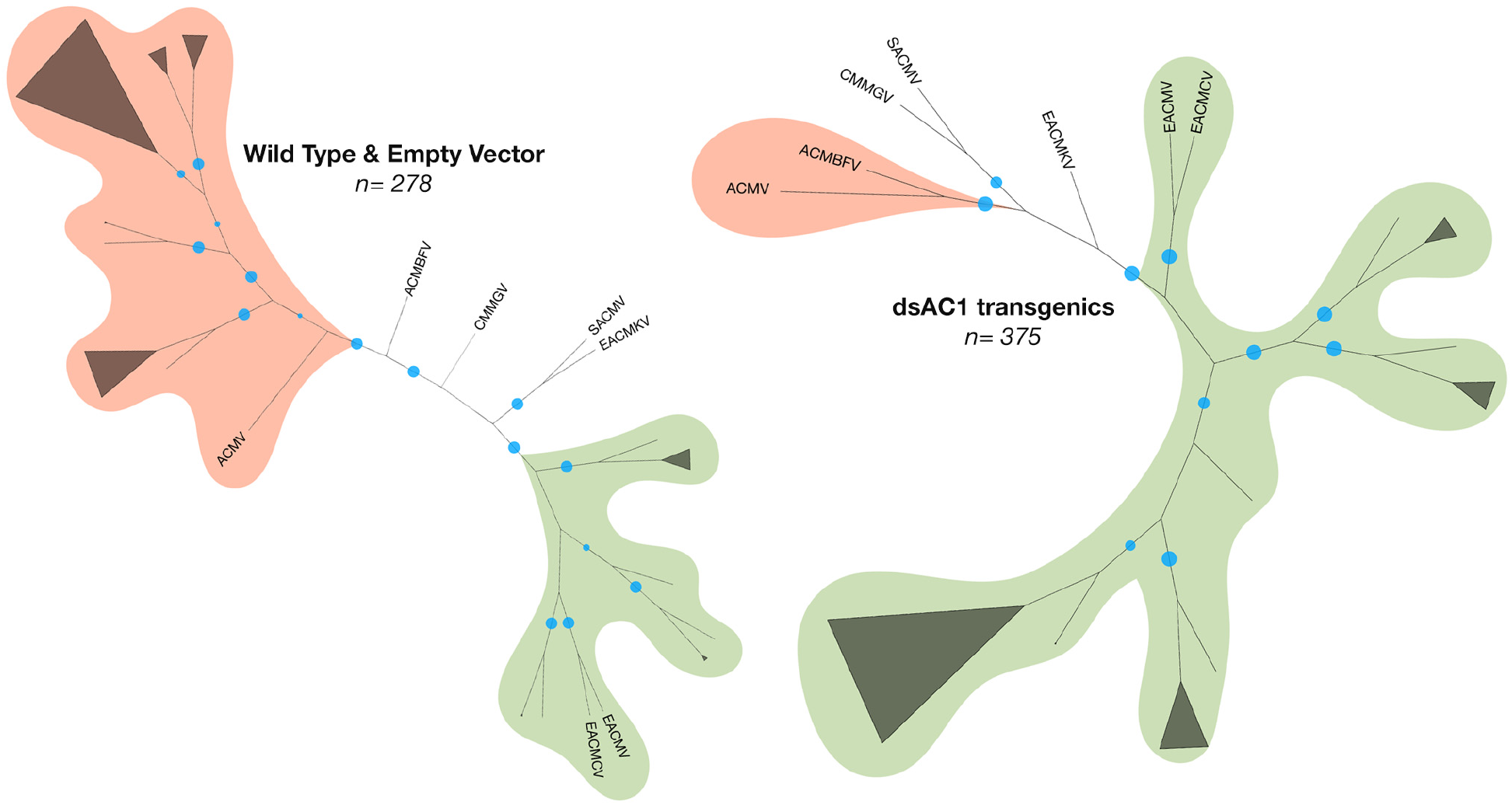
Neighbour joining trees (100 bootstraps) constructed with full length genome sequences from transgenics and control plants, with reference genomes from 7 African CMG species, rooted with the ACMV-NOg sequence. Blue circles represent nodes with >70 bootstrap support.

**Figure 7:**
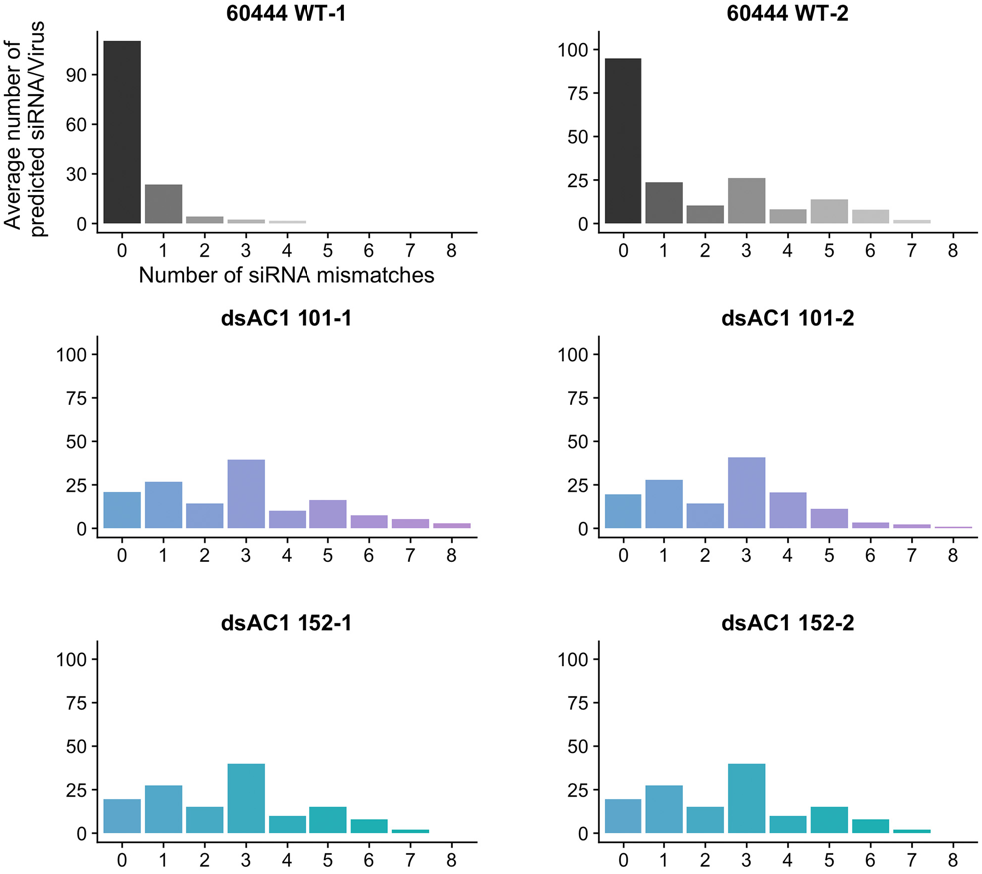
The average number of *in silico* predicted, transgene-derived 21nt siRNA which match each virus sequence obtained per sample, binned by the number of mismatches found in each case.

## DISCUSSION

In summation, CIDER-Seq effectively enriched circular DNA molecules and produced single-read, full-length DNA sequence data from the RCA reaction products. The DeConcat algorithm parsed the concatenated reads of the RCA products into individual component DNA sequences of the appropriate size range without training the algorithm with prior information of desired sequence length. Further analysis of DeConcat processing statistics provides an insight into the mechanisms of Phi29 RCA. Using infected field material, we could generate 1,328 high-quality, non-chimeric full-length virus genome sequences, representing more than twice the number of all CMG sequences deposited in GenBank to date. Moreover, according to our estimates CIDER-Seq reduces the per genome cost of sequencing by approximately 16-fold as compared to the conventional RCA and Sanger sequencing method.

RNAi technology is currently being trialed for deployment in a number of crops (Wagaba et al. 2017; Ahmed et al. 2017; Fuentes et al. 2016; Yang et al. 2017). Using CIDER-Seq we found that virus populations are radically changed in transgenic plants due to the expression of anti-viral dsRNA. We show that anti-viral cassava transgenics are able to effectively eliminate target virus species but are ineffective against species with less than 90% identity to the transgene-derived dsRNA. Thus, based on the CIDER-Seq results, we conclude that profiling virus sequence diversity is a necessary step prior to the development of new virus-resistant transgenic plants.

CIDER-Seq can also be applied to other important viruses with similar genome sizes and topology, including e.g. *Porcine circoviruses, Chicken anemia virus*, and the recently discovered, ubiquitous, human-infecting *Torque teno virus* (Ssemadaali et al. 2016). The circular dsDNA *Human papillomavirus* is another important target for future CIDER-Seq experiments. The sequencing of non-viral circular DNA templates such as plasmids is another potential application of CIDER-Seq, particularly in the context of clinical deep-sequencing of antibiotic resistance gene carrying plasmids (Conlan et al. 2014). In addition to circular DNA templates, single molecule sequencing of linear DNA and RNA viruses could also be facilitated by DeConcat in combination with size selection, Phi29 polymerase amplification and SMRT sequencing. Based on the current accuracy of long-read sequencing technologies CIDER-Seq produces results with an estimated error rate that is far below virus species and isolate demarcation thresholds. We also note that an error rate of 0.1 % is comparable to the Q30 threshold of Illumina short reads. Further improvements of either SMRT read-length or single-read quality will increase the number of full-length viral genomes and expand our method to larger viruses. Considering recent calls to incorporate metagenomics data in virus classification^1^, CIDER-Seq is a superior method for high-quality full-genome sequencing of viruses infecting plants, animals and bacteria that will facilitate building accurate sequence datasets for virus taxonomy and evolutionary studies.

## METHODS

### Field Trial Design and Planting

The field trial was conducted in an enclosed field at the Kenya Agricultural and Livestock Research Organisation site in Alupe, Kenya. The field was divided into three replicated blocks of equal size. Each block contained randomised plots of 6 plants each, per plant line (spaced at 1 m distance from each other). Plots were divided by infector plants which were multiplied from naturally infected, symptomatic plants collected from different locations in western Kenya and placed randomly around each plot. The entire field was also surrounded by healthy, non-transgenic plant rows of 2 meters in width. The study was authorised by and conducted according to the containment measures set by the National Biosafety Authority of Kenya.

Transgenic and control plants were multiplied *in vitro* and 4 week old plantlets were transferred to soil and hardened in a level II screen-house for eight weeks. Twelve week old plants were then replanted in designated positions in the field.

### Scoring and Monitoring

The field was monitored weekly for virus symptoms, whitefly populations per plant and the plant height. Symptom scoring was done according to a 5-point scale according to Ogbe et al., (2003)(Ogbe et al. 2003). CMD incidence was calculated as the proportion of infected plants in per line, per replicated block. In accordance with the biosafety guidelines, flower buds were removed and incinerated.

### Sampling and DNA extraction

Mature symptomatic cassava leaves were harvested from infected plants grown for nine months in the field. The plants included cassava genotypes 60444 and TME14 as well as transgenic lines in the same genotypic backgrounds. Total nucleic acid was extracted from leaf samples pooled from three plants of each genotype. Extraction was performed using a CTAB (cetyl trimethylammonium bromide) protocol(Chang et al. 1993) combined with an ethanol precipitation step. Total nucleic acid was quantified using a Qubit dsDNA BR Assay Kit (Q32850, Thermo Fisher Scientific).

### Size Selection

For the pre-enrichment size-selection step, 5 μg of total nucleic acid was loaded on a 0.75% agarose gel cassette and separated on a BluePippin instrument (SAGE Science). DNA fragments between 0.8-5kb were extracted. This size range was selected because geminivirus DNA is present as dsDNA replicative intermediates (running between 2 and 3 kb) and ssDNA mature forms, which migrate at lower size ranges on agarose gels. Post-enrichment size-selection was similarly performed and fragments >3 kb were extracted. An additional sequencing library was produced without the post-enrichment size selection step. This library is referred to as NSS in the following text.

### Random Circle Amplification

Random rolling circle amplification was performed as previously described (Dean et al. 2001) with some modifications. A 20 μl reaction was set up using 5 μl of size-selected template DNA, 1 mM dNTPs, 10U Phi29 DNA polymerase (EP0092, Thermo Fisher Scientific), 50 μM Exo-resistant random primer (SO181, Thermo Fisher Scientific), 0.02U inorganic pyrophosphatase (EF0221, Thermo Fisher Scientific) and 1X Phi29 DNA polymerase buffer (supplied with enzyme). The reaction was run at 30 °C for 18 hours and stopped by heating to 65 °C for 2 minutes. Product DNA was purified by sodium acetate/ethanol precipitation. We also used the illustra TempliPhi 100 amplification kit (25640010, GE Life Sciences) and obtained similar amplification results.

### Phi29 debranching

10 μg of amplified DNA was used in a debranching reaction with 5U of Phi29 DNA polymerase without a primer at 30 °C for 2 hours and stopped by heating at 65 °C for 2 minutes. The product was precipitated with sodium acetate/ethanol. The purified product was treated with 50U S1 nuclease (EN0321, Thermo Fisher Scientific) in a 20 μl reaction at 37 °C for 30 minutes and stopped by adding 3.3 μl of 0.5M EDTA and heating at 70 °C for 10 minutes. DNA was purified by sodium acetate/ethanol precipitation.

### DNA repair

De-branched DNA was treated with 3U T4 DNA polymerase (M0203L, New England Biolabs) and 10U *E. coli* DNA polymerase I (M0209L, New England Biolabs) with 1X NEBuffer 2 and 1 mM dNTPs in a 50 μl reaction. The reaction was incubated at 25 °C for 1 hour and stopped by heating at 75 °C for 20 minutes. After cooling, 5U of Alkaline Phosphatase (EF0651, Thermo Fisher Scientific) was added. De-phosphorylation was conducted at 37 °C for 10 minutes and stopped by heating to 75 °C for 5 minutes. The repaired DNA was purified using KAPA Pure Beads (KK8000, Kapa Biosystems) at a 1.5X volumetric ratio and quantified using a Qubit dsDNA BR Assay Kit (32850, Thermo Fisher Scientific).

### Semi-quantitative PCR

Semi-quantitative PCR was performed using the primers described in Supplementary Table 2. DreamTaq polymerase (EP0705, Thermo Fisher Scientific) was used to amplify 10 ng of template in 50 μl reactions set for 15, 25 and 40 cycles each. PCR products were separated using a 1 % agarose gel in 1X sodium borate acetate buffer and visualised by staining with ethidium iodide.

### SMRT barcoding, library preparation and sequencing

A Bioanalyzer 2100 12K DNA Chip assay (5067-1508, Agilent) was used to assess fragment size distribution of the enriched DNA samples. The sequencing libraries were produced using the SMRTBell™ Barcoded Adapter Complete Prep Kit - 96, following manufacturer’s instructions (100-514-900. Pacific Biosciences). Approximately 200 ng of each DNA sample was end-repaired using T4 DNA Polymerase and T4 Polynucleotide Kinase according to the protocol supplied by Pacific Biosciences. A PacBio barcoded adapter was added to each sample via a blunt end ligation reaction. The 9 samples were then pooled together and treated with exonucleases in order to create a SMRT bell template. A Blue Pippin device (Sage Science) was used to size select one aliquot of each barcoded library to enrich the larger fragments >3 kb. Both the non-size selected and the size selected library fractions were quality inspected and quantified on the Agilent Bioanalyzer 12Kb DNA Chip and on a Qubit Fluorimeter respectively. A ready-to-Sequence SMRTBell-Polymerase Complex was created using the P6 DNA/Polymerase binding kit 2.0 (100-236-500, Pacific Biosciences) according to the manufacturer instructions. The Pacific Biosciences RS2 instrument was programmed to load and sequence the samples on 1 SMRT cell v3.0 (100-171-800, Pacific Biosciences), taking 1 movie of 360 minutes. A MagBead loading (100-133-600, Pacific Biosciences) method was chosen to improve the enrichment of longer DNA fragments. After the run, a sequencing report was generated for every cell via the SMRT portal to assess the adapter dimer contamination, sample loading efficiency, the obtained average read-length and the number of filtered sub-reads.

The organisation of samples per SMRT Cell is depicted in Supplementary Table 1.

### CIDER-Seq data analysis

Following SMRT sequencing and the generation of barcode-separated subreads (Pacific Biosciences Inc. 2015) we followed a custom data analysis method. First, we implemented the RS_ReadsOfInsert.1 program using the SMRTPipe command line utility (Pacific Biosciences) using the following filtering criteria: Minimum Predicted Accuracy= 99.9, and Minimum Read Length of Insert (in bases) = 3,000. (For the error analysis, the same analysis was repeated by changing the Minimum Predicted Accuracy to 99.5 and 99.0 respectively). Resulting high quality ROIs were binned into three categories, virus DNA A, virus DNA B or non-viral DNA based on BLAST results (expect value threshold = 1.0) against a database comprised of the full-length *East African Cassava mosaic virus* (EACMV) DNA A (AM502329) and DNA B (AM502341).

### Sequence Trimming

Next, to identify the putative viral DNA sequence start and end points in each sub-read, the binned ROIs were aligned against two modified EACMV DNA sequences (DNA A and DNA B, for sequences in their respective bins) using MUSCLE (Edgar 2004). The modified sequences consisted of: a) a three times concatenated full-length genome sequence and b) a genome sequence flanked on either side by two half sequences. This was done to simulate the linearization of the circular genome and allow for the best alignment of the generated sub-read. A 10 nt sliding window was then run from both ends of the alignment. The first window (at both sequence ends) to detect a 90% sequence identity (i.e. 9 out of 10 nucleotides in the sliding window are identical) between the read and the two modified reference sequences was designated as the start and end point of the viral sequence respectively (Fig. 2, Step 4). The sequence between these two points (called the trimmed ROI) was further analysed using DeConcat (Fig. 3).

### Sequence Annotation and Phasing

The de-concatenated ROIs were annotated using tBLASTn against virus protein reference sequences (Supplementary Sequence 1) with an e-value of 0.01. The high-scoring pairs (HSPs) were summarized and sequences with protein-annotations that had contradicting and incorrect coding strands (according to the reference annotation) or lacked the full set of geminivirus proteins were eliminated. Sequences were replaced by their reverse complements where necessary to maintain all sequences in the same strand (i.e. +strand relative to the geminivirus AV1/AV2 genes). Annotation positions were also adjusted accordingly. Next, since geminiviruses have circular genomes, for comparative analyses we phased all sequences to the same start position (minus an offset) of a selected reference protein (in our case AV1). The proteins were annotated by using the start and end codon positions derived from the tBLASTn HSPs. The results were saved in FASTA (sequences only) and GenBank (sequences with ORF annotations) formats. Frameshift errors were detected using the HSP results from tBLASTn to identify cases where the same protein annotation had a break or overlap in the alignment results. The error/sequence statistic was calculated by dividing the total number of frameshifts detected by the total number of sequences in each dataset.

### Percentage Identity Calculation

Phased sequence reads and reference sequences (also phased) were first trimmed to ensure identical start and end positions. Trimmed, phased reads were pairwise aligned to the reference sequence (either ACMV-NOg full genome, for Fig. 5 or the 146bp dsRNA sequence, for Supplementary Fig. 7) using the Pairwise2 package in BioPython. Identity scores were calculated for each pairwise alignment and plotted on a density plot using the ggjoy/ggridges R package.

### Phylogenetic Analysis

Trimmed, phased reads were aligned using MUSCLE implemented in the CLC Genomics Workbench 10.0 using default parameters. The alignment was used to create a Neighbour-Joining tree with 100 bootstraps. The phylogeny was used to create an unrooted cladogram and clades were defined based on the position of the 7 reference sequences used.

### siRNA simulation

We first divided the dsRNA complementary region of the transgene into 21 nt fragments using a 1 nt sliding window. These putative siRNAs were then locally aligned (with a high gap open/extend penalty) against each full-length trimmed, phased virus sequence read obtained (see above) per sample. The number of mismatches present in each alignment was recorded. Results were tabulated showing the number of siRNA with no mismatches, with 1 mismatch, and so on as plotted in Fig. 6. This process was implemented in Python and the code is available for download below.

## DATA ACCESS

Python packages for the CIDER-Seq Data Analysis software and DeConcat, along with installation and usage guidelines are available at: http://www.dx.doi.org/10.5281/zenodo.834928. Python scripts for siRNA counting, sequence extraction and alignment, and R scripts for plotting data are available at: www.dx.doi.org/10.5281/zenodo.1009036.

Full length annotated genome sequences, raw sequence data and data generated during intermediate CIDER-Seq and siRNA simulation steps are freely available at: www.dx.doi.org/10.5281/zenodo.830530 and at: www.dx.doi.org/10.5281/zenodo.1009036.

## ACKNOWLEDGEMENTS

This research was supported by a grant from ETH Zurich and Stiftung fiat panis to perform a confined cassava field trial in Kenya. We are grateful to the staff at the Kenya Agricultural and Livestock Research Organisation and the Masinde-Muliro University of Science and Technology, Kenya for their assistance with the field trial and the National Biosafety Authority for approvals and guidance. We thank Weihong Qi (Functional Genomics Center Zurich, Switzerland) for assistance with SMRT sequencing as well as helpful advice and discussion. Devang Mehta was supported by the European Union’s Seventh Framework Programme for research, technological development, and demonstration (EU GA-2013-608422-IDP BRIDGES).

## AUTHOR CONTRIBUTIONS

Conceptualization: DM, WG & HV; Methodology and Formal Analysis: DM, MHH; Investigation: DM, MW, HW, AP; Software: MHH; Visualisation, Writing-original draft: DM; Funding acquisition, Resources, Writing-review & editing, Supervision: WG & HV. All authors agree with the final version of the manuscript.

## SUPPLEMENTAL INFORMATION

**Supplementary Table 1:**
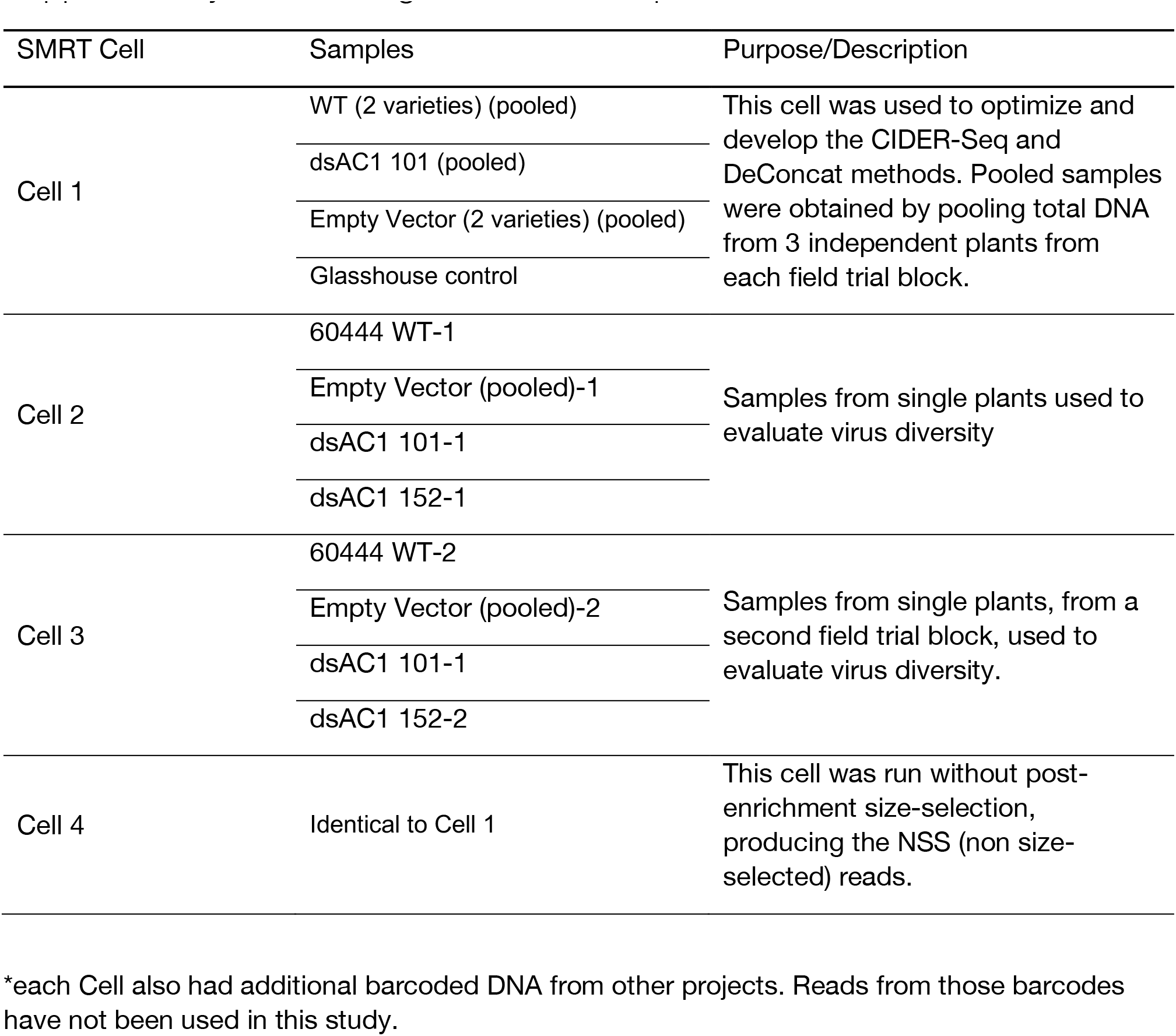
Organization of samples in SMRT Cells

| SMRT Cell | Samples | Purpose/Description |
| --- | --- | --- |
| Cell 1 | WT (2 varieties) (pooled) | This cell was used to optimize and develop the CIDER-Seq and DeConcat methods. Pooled samples were obtained by pooling total DNA from 3 independent plants from each field trial block. |
|  | dsAC1 101 (pooled) |  |
|  | Empty Vector (2 varieties) (pooled) |  |
|  | Glasshouse control |  |
| Cell 2 | 60444 WT-1 | Samples from single plants used to evaluate virus diversity |
|  | Empty Vector (pooled)-1 |  |
|  | dsAC1 101-1 |  |
|  | dsAC1 152-1 |  |
| Cell 3 | 60444 WT-2 | Samples from single plants, from a second field trial block, used to evaluate virus diversity. |
|  | Empty Vector (pooled)-2 |  |
|  | dsAC1 101-1 |  |
|  | dsAC1 152-2 |  |
| Cell 4 | Identical to Cell 1 | This cell was run without post-enrichment size-selection, producing the NSS (non size-selected) reads. |

\*each Cell also had additional barcoded DNA from other projects. Reads from those barcodes have not been used in this study.

**Supplementary Table 2:**
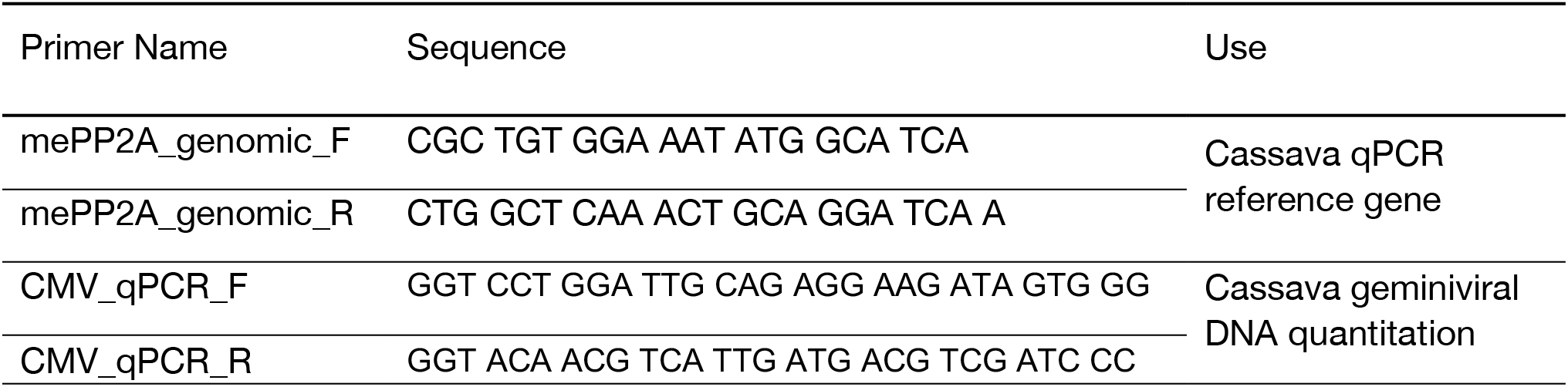
Primer sequences

**Supplementary Figure 1:**
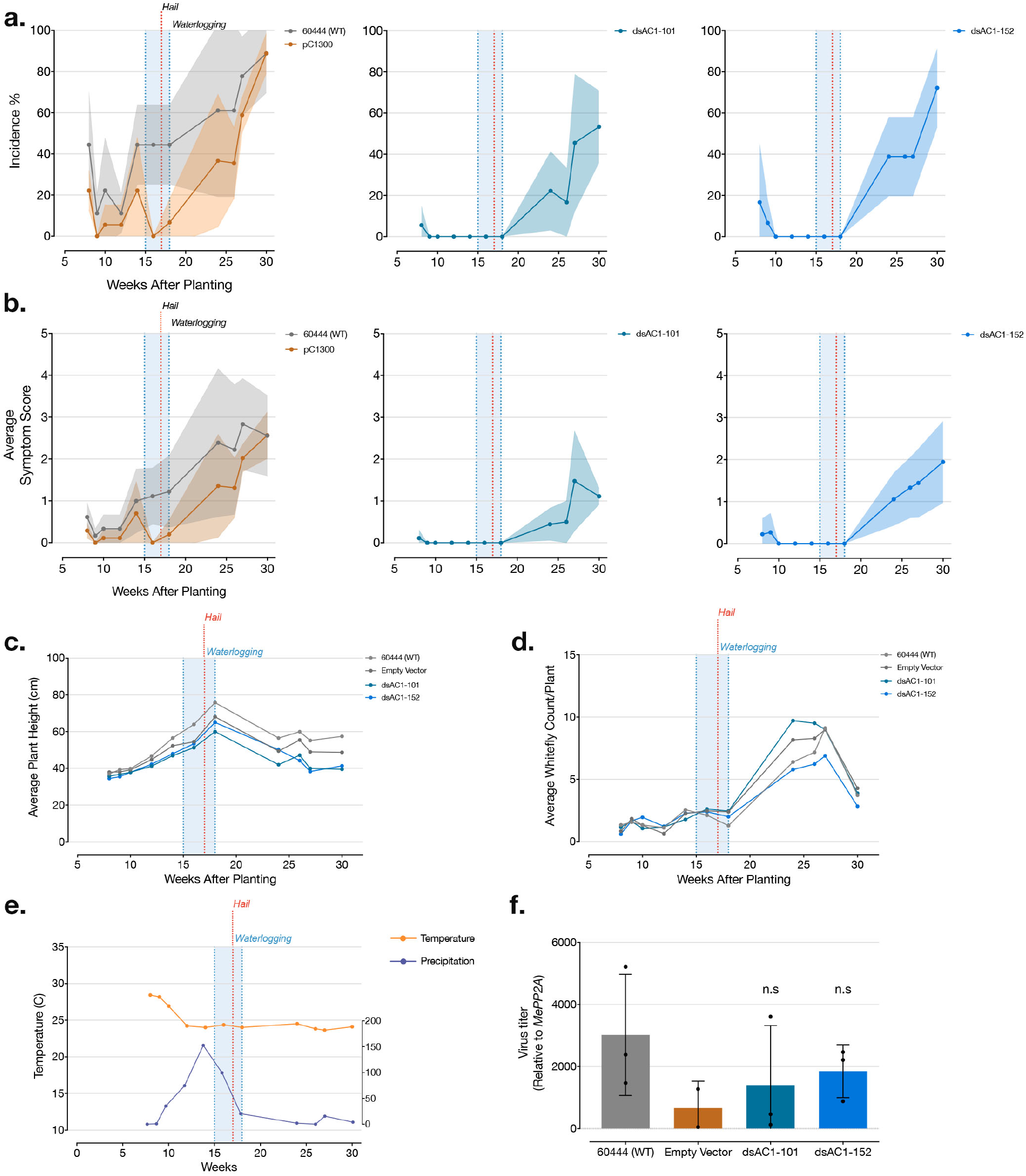
Field data. **(a)** Disease incidence (% of symptomatic plants per line). Shaded area shows the standard deviation across three replicated blocks **(b)** Average symptom severity scores per line. Shaded area shows the standard deviation across three replicated blocks. **(c)** Average plant height and **(d)** whitefly counts per plant, per line in all three blocks **(e)** Rainfall and temperature measurements at the field location. Shaded blue are shows three weeks of waterlogging at the field site and the red dotted line indicates the occurrence of a hailstorm **(f)** Virus titres in pooled samples per line, from each replicated block measured by quantitative PCR relative to plant genomic DNA.

**Supplementary Figure 2:**
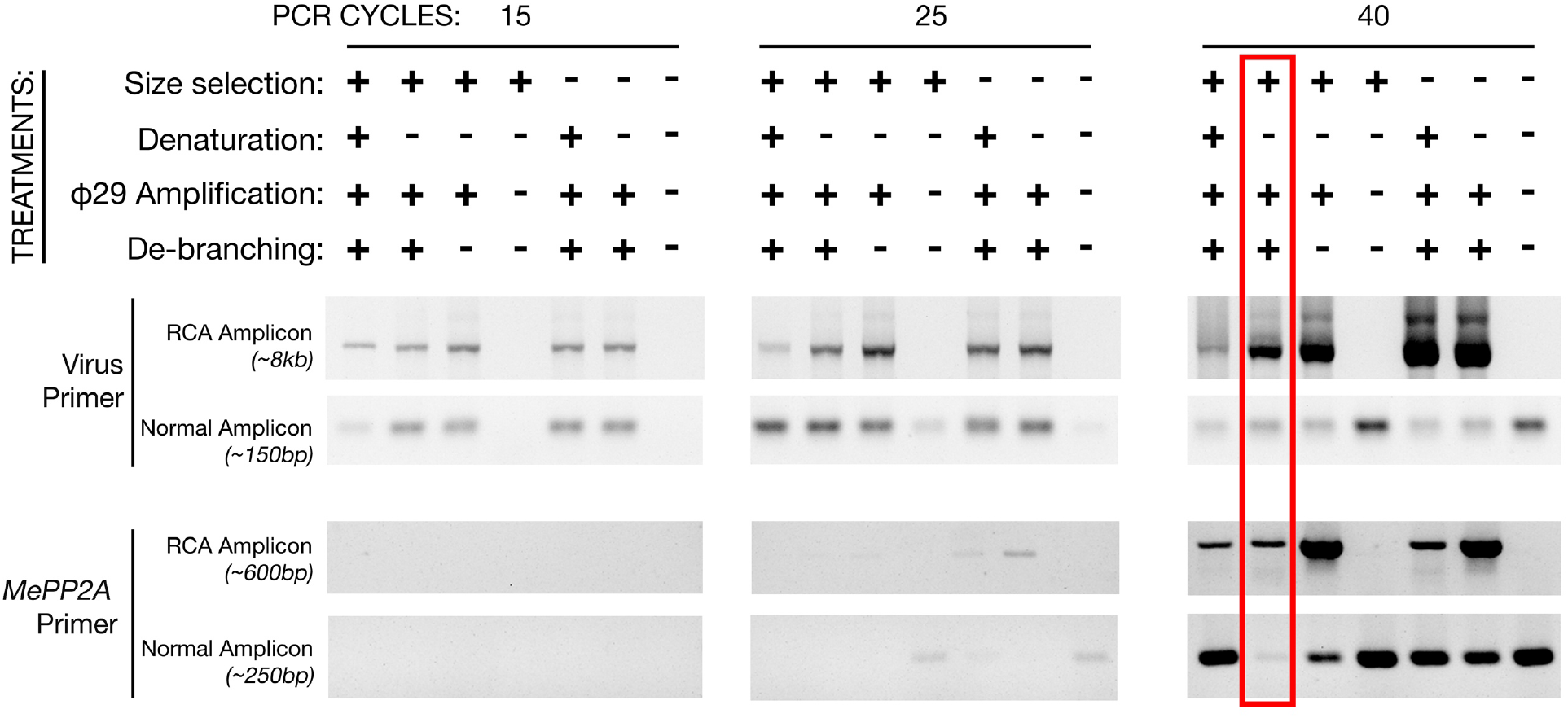
Semi-quantitative PCR on a sample testing permutations of the enrichment steps shown in Figure 1. *Manihot esculenta PROTEIN PHOSPHATASE 2A (MePP2A)-* specific primers were used to determine the amount of linear cassava genomic DNA compared to enriched viral circular DNA amplified with cassava geminivirus-specific primers. The result from the selected enrichment protocol is highlighted in red.

**Supplementary Figure 3:**
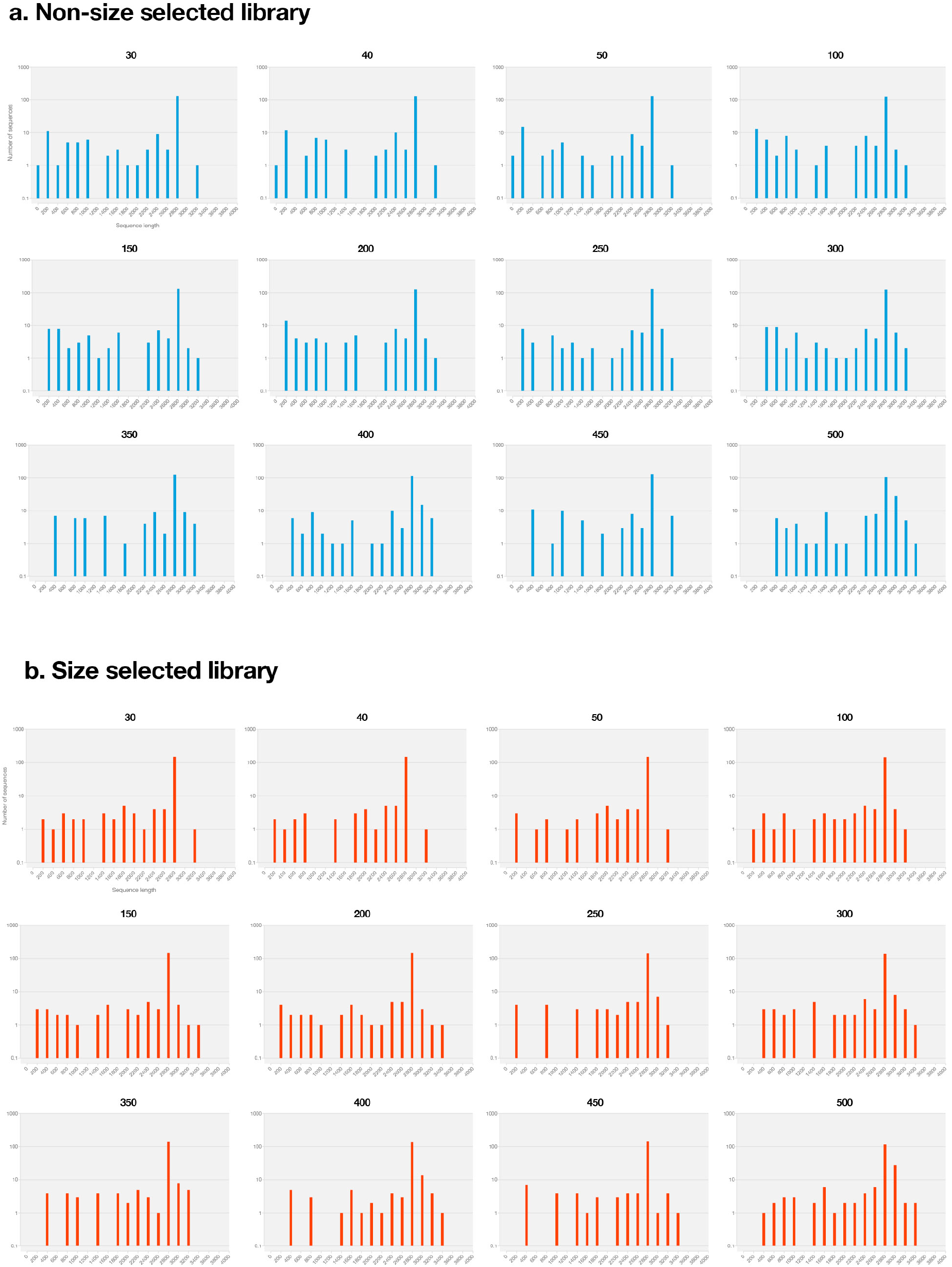
DeConcat benchmarking with differently sized cuts in step 1. **(a)** non-size selected library, **(b)** size-selected library (>3 kb). (Data from Cell 1 and 4, Supplementary Table 1)

**Supplementary Figure 4:**
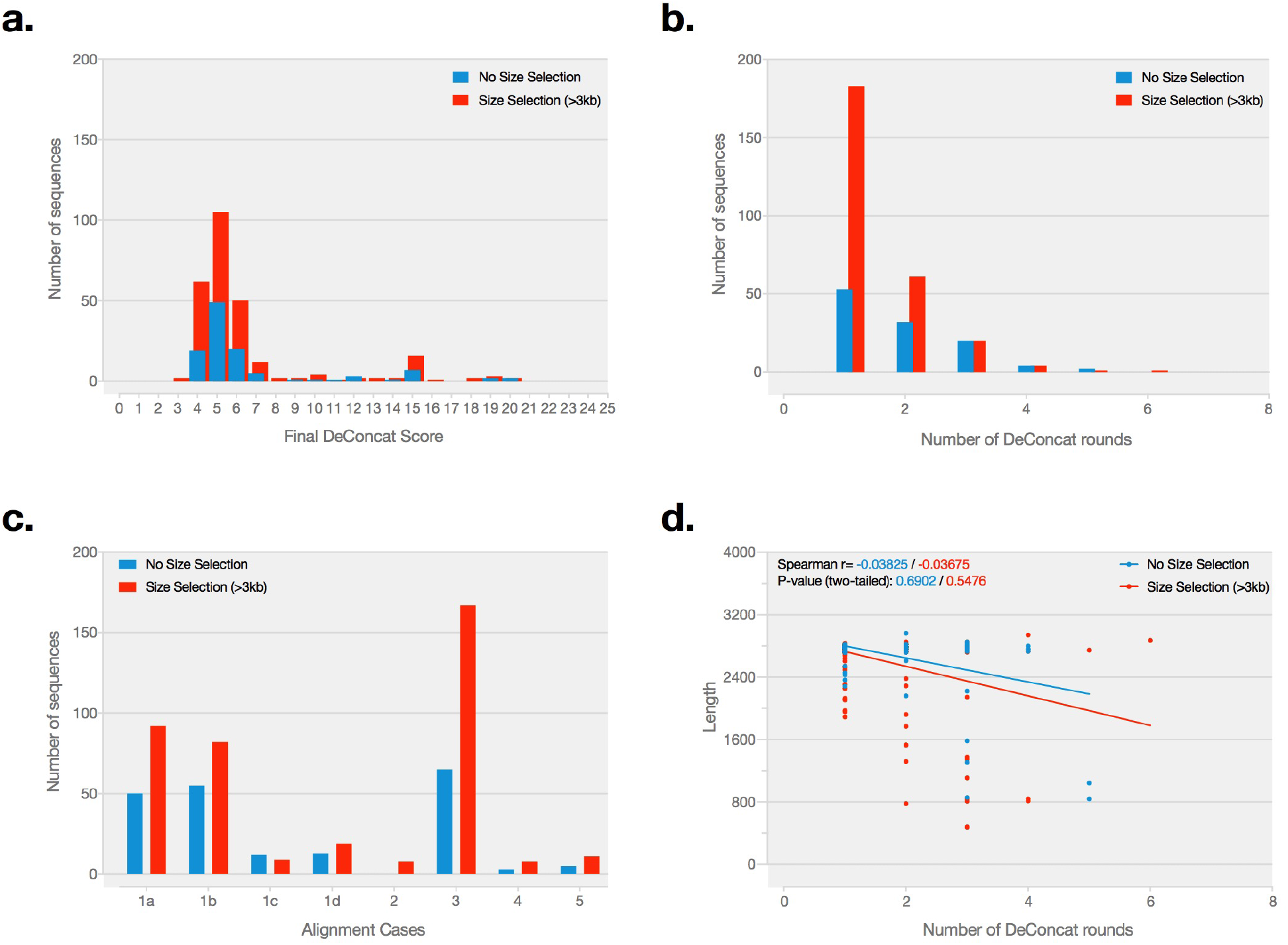
Performance of the DeConcat algorithm **(a)** Frequency curve of the number of DeConcat rounds required to process each sequencing read. **(b)** Frequency curve of the final DeConcat alignment score obtained for the CMG datasets. **(c)** Frequency of different alignment forms used during DeConcat. **(d)** Correlation plot of the number of DeConcat rounds required to completely resolve sequences versus the length of the final de-concatenated sequence.

**Supplementary Figure 5:**
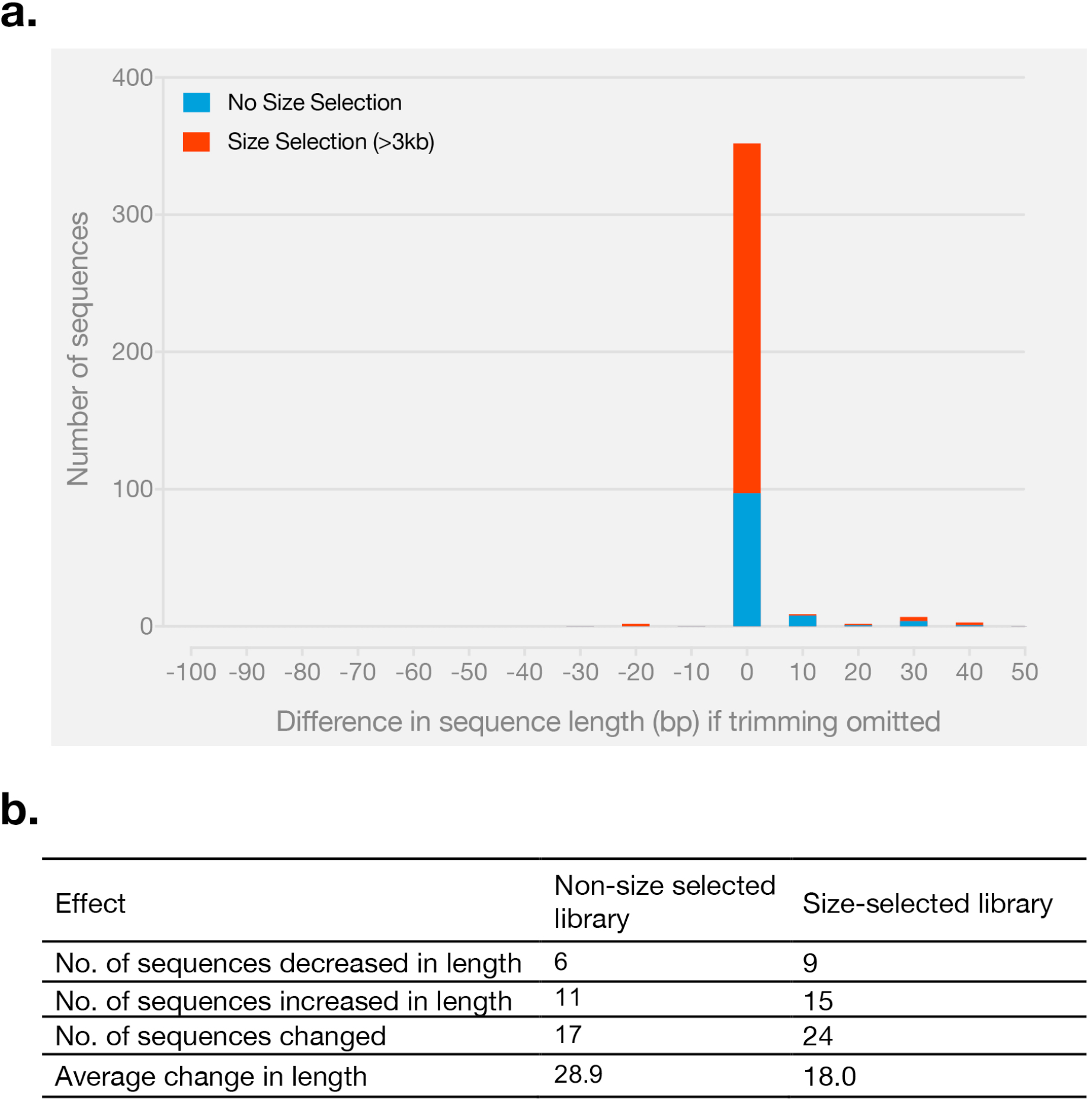
The effect of applying DeConcat directly on sequencing reads without reference-based trimming. **(a)** Number of sequences (y-axis) versus the change in bp (x-axis) without trimming. **(b)** Summary of changes without trimming.

**Supplementary Figure 6:**
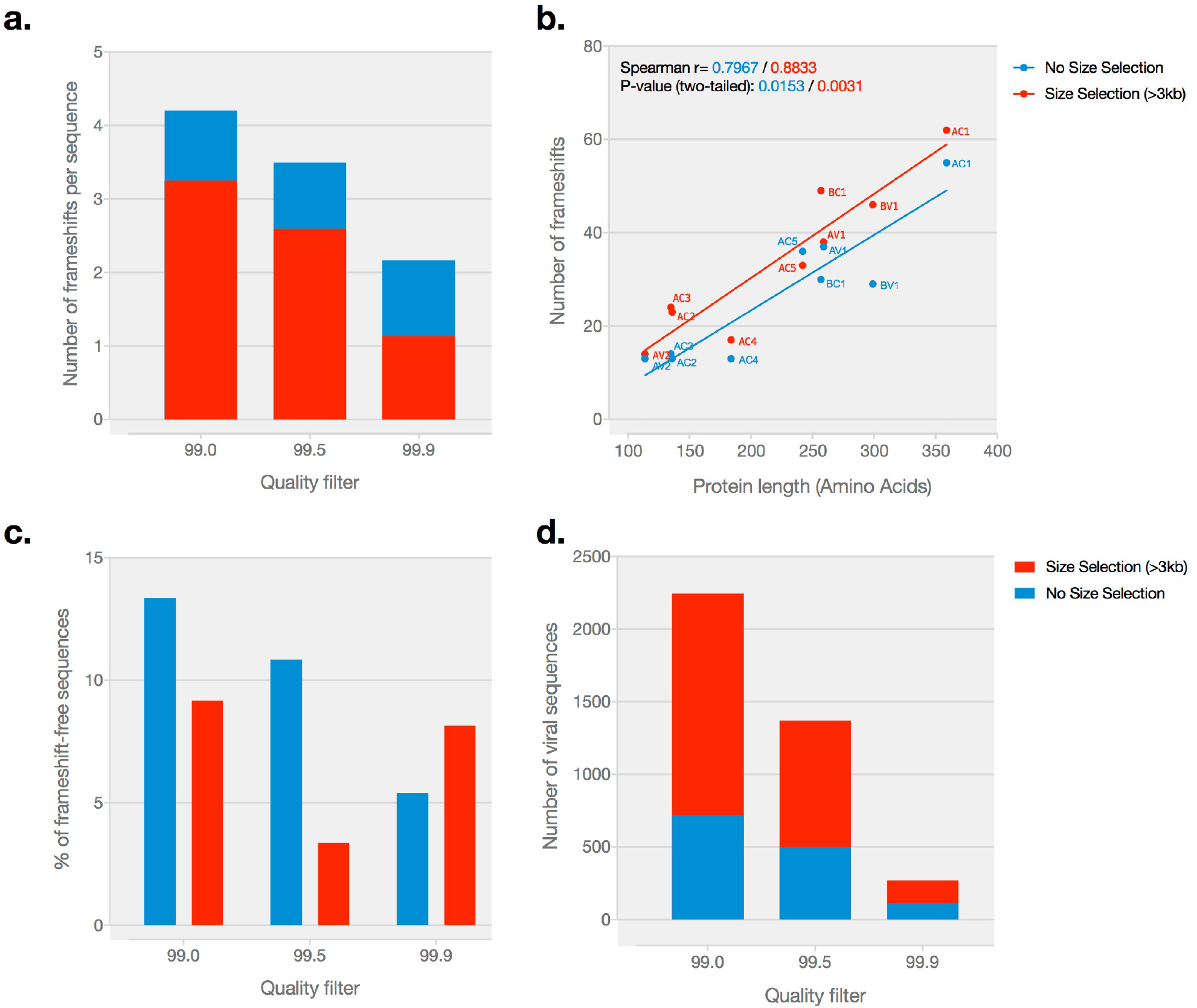
Analysis of frameshift mutations in de-concatenated CMG sequences using different SMRT sequence quality thresholds. **(a)** The number of frameshifts decreases with increasing quality thresholds. **(b)** Number of frameshifts per protein correlates with the amino acid sequence length of the respective protein. **(c)** Percentage of frameshift-free sequences. Performance of the DeConcat algorithm. **(d)** Total number of CMG sequences produced using different quality thresholds for SMRT analysis. (Data from Cell 1 and Cell 4, Supplementary Table 1)

**Supplementary Figure 7:**
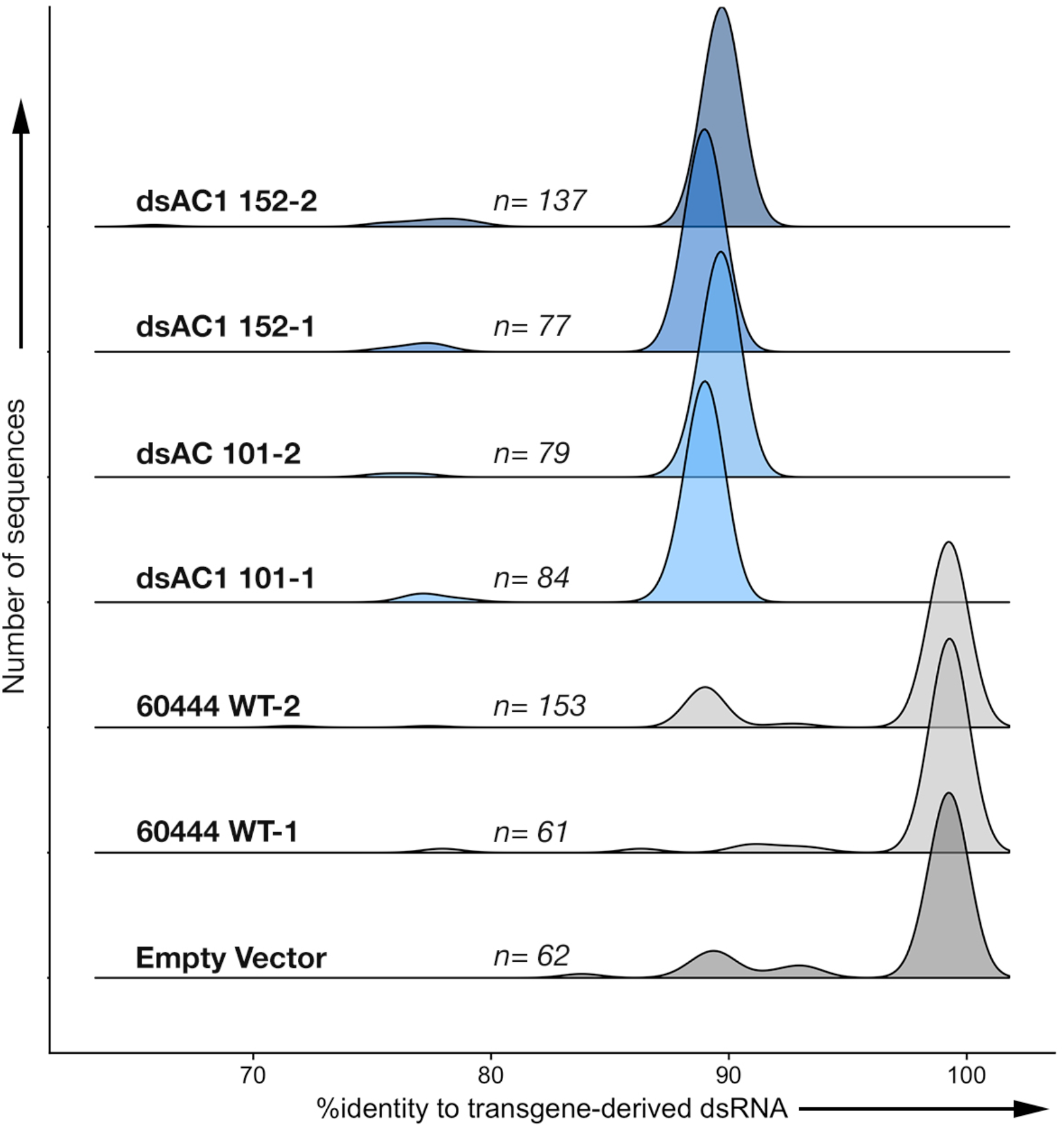
Density plot showing the distribution of virus sequences in each sample with their %identity to the transgene-derived dsRNA sequence.

**Supplementary Sequence 1:** Reference protein sequences from the *East African cassava mosaic virus*.

>AV2

MWDPLVNEFPDSVHGLRCMLAIKYLQALEDTYEPSTLGHDLVRDLVSVIRARNYVEATRRYHHFHSRLEGSSKAELRQPIQEPCYCPHCPRHKSKTGLDKQAHVQKAHNVQDV*

>AV1

MSKRPGDIIISTPGSKVRRRLNFDSPYRNRATAPTVHVTNRKRAWINRPMYRKPTMYRMYRSPDIPRGCEGPCKVQSYEQRDDVKHLGICKVISDVTRGPGLTHRVGKRFCIKSIYILGKIWMDENIKKQNHTNNVMFYLLRDRRPYGNAPQDFGQIFNMFDNEPSTATIKNDLRDRFQVLRKFHATVIGGPSGMKEQALVKRFYRLNHHVTYNHQEAGKYENHTENALLLYMACTHASNPVYATLKIRIYFYDSIGN*

>AC4

MPFRDTYIMSPINRRRCSRVSLVECLQLTWSTCELRVLEFKPRMSFSHTQSVLYPKNTSCHSFKHYLSHQTLSSLKSVESCIRMGNLTCMPSSNSRGKSRLRTIVSSIVYTQPVAPISTPTFKVPNQAQMSSPIWIRTETPSNGDNFRSMDDLLEAVNNQRMMLTPKALTAAVSQKLLMSLGN*

>AC1

MRTPRFRVQAKNVFLTYPKCSIPKEHQLSFIQTLSLPSNPKFIKICRELHQNGEPHLHALIQFEGKITITNNRLFDCVHPTCSTNFHPNIQGAKSSSDVKSYLDKDGDTVEWGQFQIDGRSARGGQQSANDAYAKGLNSGSKSEALNVIRELVPKDFVLQFHNLNSNLDRIFQEPPAPYVSPFPCSSFDQVPVEIEEWVADNVRDSAARPWRPNSIVIEGASRTGKTIWARSLGPHNYLCGHLDLSPKVFNNAAWYNVIDDVDPHYLKHFKEFMGSQRDWQSNTKYGKPVQIKGGIPTIFLCNPGPTSSYKEFLDEEKQEALKAWALKNAIFITLTEPLYSGSNQSQSQTIQEASHPA*

>AC2

MQSSSPSQNHSTQVPIKVSHRQFKKRAIRRRRVDLVCGCSYYLHINCSNHGFTHRGTHHCSSSNEWRVYLGNKQSPVFHNHQAPTTTIPAEPGHHNSPGSIQSQPEEGAGDSQMFSQLQDLDDLTASDWSFLKGL*

>AC3

MDLRTGELITAPQAMNGVYTWEINNPLYFTITRHQQRPFLLNQDIITVQVRFNHNLRKELGIHKCFLNFRIWTTLRPQTGLFLRVFRYQVLKYLDNIGVISINDVIRAADHVLFNVIAKTIECQLTHEIKFNVY*

>AC5

MSTCHVQKQSILCVILILPCLLMIICHVMIQPVKPFNQSLLLHARWTTNNSGMKFPQYLKPIPQIVLNCCSTGLIIKHIKYLPKVLGRIAIRPSIPKQIKHHIIRVILLLNIFIHPDLTKYVNGLDTKPLSDPVCQPRPTCHITNHLTDTKVLHIIPLLIRLDLTWAFTAPRYVWASIHPVHCGLSVHGPVYPGPFSICDVDSGGSSTVPVWAVEVQPSTNLRAWSGNDDISWSLRHNYEP*

>BV1

MYSIRKQPRNFQRKCNSNTTNRFPIRRKYVGGHTRPSVRRRLSYEPVERPLVYNVLCEKQHGDVFNLQQNTSYTSFVTYPSRGPSGDGRSRDYIKLQSMSVSGVIHAKANCNDDPMEVSHVVNGVFVFSLIMDTKPYLPAGVQALPTFEELFGSYSASYVNLRLLNNQQHRYRVLHSVKRFVSSAGDTKVSQFRFTKRLSTRRYNIWASFHDGDLVNAGGNYRNISKNAILVSYAFVSEHSMSCKPFVQIETSYVG*

>BC1

MDTSVPVISSDYIQSARTEYKLTNDESPITLQFPSTIERTRVRIMGKCMKVDHVVIEYRNQVPFNAQGSVIV

TIRDTRLSDEQQDQAQFTFPIGCNVDLHYFSASYFSIDDNVPWQLLYKVEDSNVKNGVTFAQIKAKLKLSAA

KHSTDIRFKQPTIKILSKDYGPDCVDFWSVGKPKPIRRLIQNEPGTDYDTGPRYRPITVQPGETWATKSTIG

RSTSMRYTGPKHIDIDDSSSKQYASEAEFPLRGLHQLPEASLDPGDSVSQTQSMSKKDIESIIEQTVNKCLI

AHRGSSHKDL*

